# Ergodic breakdown of brain dynamics under chronnectome complexity across cognitive states and age

**DOI:** 10.64898/2026.11.06.764964

**Authors:** Behnam Kazemivash, David C. Zhu

## Abstract

Chronnectome complexity, the extent to which time-varying functional connectivity follows a structured sequence of distinct configurations, contributes to shaping large-scale brain dynamics. However, it remains unclear how this complexity influences the encoding of information that distinguishes resting state and task conditions. In ergodic theory, time averages represent ensemble behavior under ergodicity, which breaks down when dynamics are non-stationary, as when activity moves sequentially between distinct regimes. We hypothesize that this information resides in time-averaged connectivity under lower-complexity conditions but in trajectory geometry under higher-complexity conditions, where brain activity traverses distinct configurations over time, consistent with ergodic breakdown, which we term the *information migration principle*. We formulate this as dynamics-driven multi-class classification and introduce the Brain Dynamics Classifier (BDC), an interpretable data-driven mechanistic model in which a graph neural network embeds dynamic functional connectivity into a latent space and a neural ordinary differential equation learns the vector field governing trajectories through that space. The proposed model derives two complementary representations within a single learned dynamical system, capturing time-averaged connectivity, which is sufficient under ergodic dynamics, and trajectory geometry, which preserves the temporal organization obscured by averaging under non-ergodic dynamics. Our results show high classification accuracy, **97.4%** for eight conditions in the Human Connectome Project young adult cohort and **92.2%** for four in the aging cohort. To test the information migration principle, we ablated the time-averaged representation and found that trajectory contributions varied with chronnectome complexity, from high for Working Memory to absent for Relational Reasoning, consistent with ergodic breakdown. Including subcortical regions further increased trajectory contributions for affective conditions. In aging, the distinction between higher and lower complexity conditions was preserved, resting-state dynamics became less ergodic with advancing age, and higher fluid intelligence was associated with more ergodic resting-state dynamics.

## 1 Introduction

The human brain is a complex dynamical system in which cognitive function emerges not only from anatomical connectivity but also from the time-varying coordination of activity across distributed networks [1–3]. These networks reconfigure across the sequential phases of complex tasks, the encoding and retrieval stages of memory, and sustained attentional control [4–6]. The temporal organization of connectivity, known as the chronnectome [7], carries information about cognitive state beyond time-averaged summaries [8]. However, it has been characterized mainly through descriptive methods that summarize which connectivity states occur and how they change, without modeling the rules that govern their evolution or identifying which aspects of this organization carry information about cognitive state [9, 10]. It therefore remains unclear *where* condition-discriminative information resides, in time-averaged connectivity or in the trajectory geometry of connectivity. A candidate determinant is chronnectome complexity, the complexity of time-varying connectivity patterns, which we define as the extent to which a condition imposes a temporally structured sequence of distinct connectivity configurations rather than sustaining a single configuration. Chronnectome complexity is a property of brain dynamics rather than of task design, so we quantify it from the learned dynamics instead of imposing an a priori ordering. It also differs from task difficulty and from the breadth of dynamics, because a task can be computationally demanding yet unfold within a largely stable configuration, and a state such as rest can explore many configurations without an imposed sequence. Whether the locus of this information shifts systematically with chronnectome complexity has direct consequences for how cognitive states are measured and for downstream analyses of brain function across the lifespan.

Attractor landscape theory provides a conceptual foundation for this question, describing cognition as the traversal of an energy landscape whose geometry determines whether brain activity settles into a stable configuration or moves continuously between functionally distinct states [11–13], consistent with evidence for metastable switching and graded transitions between segregated and integrated functional architectures [3, 14, 15]. Ergodic theory formalizes this distinction. A dynamical system is ergodic when its time averages equal its ensemble averages [16, 17]. When brain activity remains near a stable attractor, or repeatedly revisits a common attractor repertoire, its statistics remain stationary and its time average faithfully represents brain state. In contrast, when activity traverses distinct configurations in a sequence that diverges from these configurations, the time average becomes a *ghost state*, a superposition of configurations visited sequentially but never occupied simultaneously. Unlike the ghost attractors of multistable network models [12], which are latent dynamical states, the ghost state is a statistical artifact of averaging over a non-stationary sequence. In this case, condition-discriminative information should reside in the temporal geometry of the trajectory rather than in its mean. We formalize this hypothesis as the *information migration principle*, under which the locus of condition-discriminative information shifts with chronnectome complexity from time-averaged connectivity under lower-complexity conditions to the geometry of continuous latent trajectories under higher-complexity conditions.

An empirical test of this principle exposes a methodological gap beyond the common descriptive approaches. Deep learning models of dynamic functional connectivity, including spatio-temporal graph networks and transformers, classify cognitive states with high accuracy [18–20] but learn entangled representations in which time-averaged and dynamic information cannot be separated, whereas latent dynamical models such as LFADS and neural ODEs learn explicit continuous dynamics [21–23] but have mainly been used to reconstruct or forecast activity rather than to identify which aspects of the dynamics distinguish cognitive states. We address this gap by formulating the problem as dynamics-driven multi-class classification, in which each dynamic functional connectivity sequence is assigned to its resting-state or task condition, and introducing the Brain Dynamics Classifier (BDC), a data-driven mechanistic and interpretable model within NeuroGraphDynamics (NGD), a modular framework for learning continuous brain dynamics. The model combines a graph attention network [24] that embeds each connectivity state into a latent space with a neural ordinary differential equation [22] that learns the vector field governing trajectories through that space. From this single learned dynamical system, it derives time-averaged and trajectory representations that enter the classifier separately, so that ablating either at inference quantifies its contribution to each condition without confounding it with differences in architecture or training. The distance between the mean of the embedding sequence and the mean of the learned trajectory, which we term the *ergodicity gap*, compares the time average of the observed connectivity states with that of a path generated by the learned dynamics, and thus reflects how strongly the observed temporal structure constrains those dynamics. It thereby provides a model-derived index of ergodic breakdown and a quantitative measure of chronnectome complexity, with smaller gaps indicating less ergodic, higher-complexity dynamics.

We apply this framework to healthy young adult and aging cohorts from the Human Connectome Project [25, 26] under cortical and cortico-subcortical parcellations, and find that where condition-discriminative information resides depends on chronnectome complexity. This work makes three contributions. First, it proposes the *information migration principle*, which grounds the location of condition-discriminative information in ergodic theory and links it to chronnectome complexity. Second, it formulates this principle as a dynamics-driven multi-class classification problem and introduces a data-driven mechanistic model that derives time-averaged and trajectory representations from a single learned dynamical system, together with the ergodicity gap as a model-derived measure of chronnectome complexity. Third, it uses this framework to show how subcortical regions and aging shape where this information resides. Our results support a view of brain dynamics in which the locus of condition-discriminative information is identified by modeling how connectivity evolves.

## 2 Method

The Brain Dynamics Classifier (BDC), implemented within the NeuroGraphDynamics framework, is a data-driven mechanistic model that combines a graph attention network for spatial connectivity encoding, a neural ordinary differential equation (neural ODE) as the deterministic dynamical engine, and a trajectory feature extractor and classification head. Given a dynamic functional connectivity (dFC) sequence and its label, where the classes are resting state and the individual task conditions, the model learns a single latent dynamical system from which two representations are derived, a time-averaged representation and a trajectory representation. The model is trained end-to-end to classify resting-state and task conditions from dFC sequences. At inference, either representation is ablated in turn to assess its contribution to condition classification. The ergodicity gap is also computed from the embedding and trajectory sequences to quantify chronnectome complexity. Together, these analyses provide a test of the information migration principle. The framework operates in two stages. First, a module computes dFC from preprocessed fMRI data and stores the resulting sequences. The model is then trained on these stored sequences. Both stages are described in the following subsections, and Figure 1 provides an overview.

**Fig. 1.**
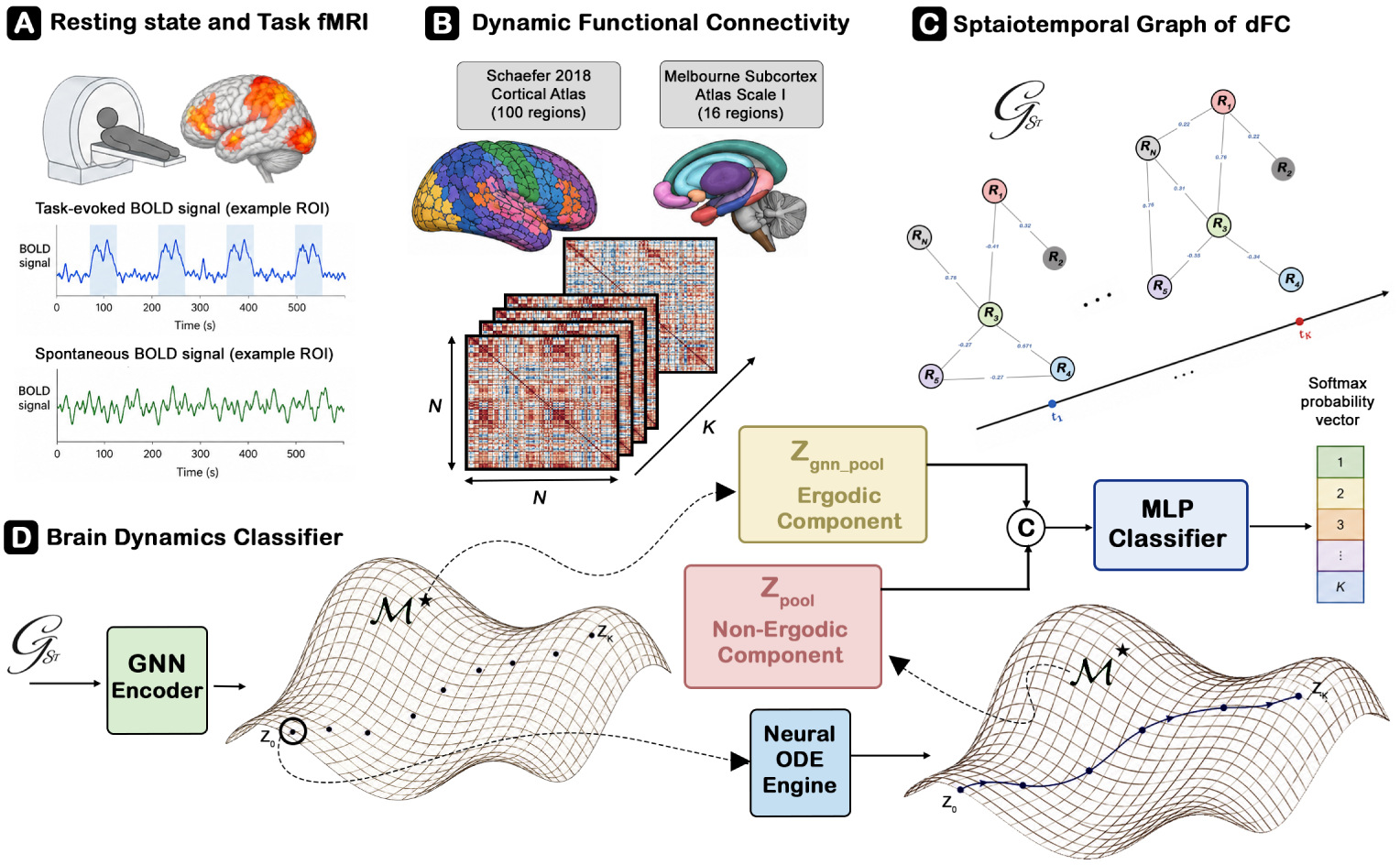
Overview of the NeuroGraphDynamics framework. (**a**) Resting-state and task fMRI data from HCP datasets. (**b**) Dynamic functional connectivity is estimated by sliding-window Pearson correlation of parcellated BOLD timeseries, using the Schaefer 2018 cortical parcellation (100 ROIs), optionally extended with Scale I of the Melbourne Subcortex Atlas (16 ROIs), yielding a dFC sequence 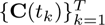. (**c**) Each dFC matrix is represented as a graph with ROIs as nodes and functional connectivity weights as edges, retaining edges with *|C_ij_ | ≥* 0.2. (**d**) In the Brain Dynamics Classifier (BDC), a GATv2 spatial encoder maps each graph to a graph-level latent embedding. A neural ODE, initialized from the first embedding, evolves a continuous latent trajectory **Z**_traj_ under a learned vector field. Trajectory features and the projected temporal mean of **Z**_gnn_ are concatenated and passed to an MLP classifier that predicts the condition.

### 2.1 Datasets and Preprocessing

We used fMRI data from two Human Connectome Project cohorts. HCP Young Adult (HCP-YA) [25] was acquired on a 3T Siemens Skyra (TR = 720 ms, voxel size = 2 mm isotropic) and covers seven task paradigms, emotional face matching (EMOTION), reward-based decision making (GAMBLING), story versus math comprehension (LANGUAGE), movement execution (MOTOR), relational reasoning (RELATIONAL), social cognition (SOCIAL), and working memory (WM), each yielding one task condition, which together with resting-state (REST) produce eight conditions. HCP Aging (HCP-Aging) [26], obtained from the Aging Adult Brain Connectome (AABC) Release 1 baseline visits, was acquired on a 3T Siemens Prisma (TR = 800 ms, voxel size = 2 mm isotropic) and covers go/no-go response inhibition (CARIT), associative face-name memory encoding (FACENAME), and visually guided motor tracking (VISMOTOR), yielding four conditions together with REST. Minimally preprocessed, ICA-FIX [27] denoised data from the HCP pipelines [28] were used without additional filtering or nuisance regression, as ICA-FIX denoising addresses structured noise including motion artifacts prior to parcellation. Subjects with missing or incomplete runs were excluded during quality control, yielding the final samples in Table 1.

**Table 1.** Participant demographics and cohort characteristics.

| **Cohort** | **Subjects<br>(train/test)** | **Age (years)** | **Gender (F/M)** | **Condition Labels** |
|---|---|---|---|---|
| HCP-YA | 981<br>(784/197) | 28.7 ± 3.7<br>(22–37) | 526/455 | 8<br>(7 tasks + REST) |
| HCP-Aging | 1,122<br>(897/225) | 61.5 ± 15.9<br>(36–90) | 639/483 | 4<br>(3 tasks + REST) |

### 2.2 Brain Parcellation and Dynamic Functional Connectivity

We evaluated two atlas configurations to probe the boundaries of the cortical observability window. The cortical atlas comprised 100 brain regions from the Schaefer 2018 parcellation [29] with 17-network assignments, spanning the default mode (Default A/B/C), executive control (Control A/B/C), somatomotor (SomMot A/B), visual (central and peripheral), limbic (Limbic A/B), salience and ventral attention (SalVentAttn A/B), dorsal attention (DorsAttn A/B), and temporoparietal (TempPar) networks. The cortico-subcortical atlas extended this with 16 subcortical regions derived from Scale I of the Melbourne Subcortex Atlas [30], including bilateral amygdala, hippocampus, anterior and posterior thalamus, caudate, putamen, nucleus accumbens, and globus pallidus, yielding 116 regions in total. The cortical-atlas experiments establish the primary results within a well-characterized cortical network parcellation, and the cortico-subcortical experiments test whether trajectory contributions for specific conditions depend on the inclusion of subcortical circuits. Mean BOLD signal within each region was extracted from the volumetric data in MNI space using the atlas label maps. Dynamic functional connectivity was estimated using sliding-window Pearson correlation [9, 31]. The parcellated BOLD timeseries **X** ∈ ℝ*^T ×N^* was segmented into overlapping windows of length *W* = 30 with a step size of *S* = 1. For each window centered at time *t_k_*, **C**(*t_k_*) ∈ ℝ*^N×N^* denotes the Pearson correlation matrix of the *W* observations within that window, with entry *C_ij_*(*t_k_*) giving the correlation between regions *i* and *j*. This yields a sequence 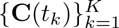 length *K* = ⌊(*T* −*W*)*/S*⌋+1, which serves as the input to the spatial encoder described in the following subsection. To remove sequence length as a confound for condition identity, the first *T*_max_ windows of each sequence were retained, with *T*_max_ = 147 for HCP-YA and *T*_max_ = 155 for HCP-Aging.

### 2.3 Model Architecture

Given a dFC sequence 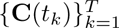 of length *T* = *T*_max_, BDC assigns a condition label *y* ∈ {1*,…, M*}, with *M* = 8 for HCP-YA and *M* = 4 for HCP-Aging. BDC comprises four components, a spatial encoder that captures *what* the connectivity state is at each window, a neural ODE that captures *how* these states evolve continuously in time from the encoded initial state, a trajectory feature extractor, and a classification head.

#### 2.3.1 Spatial Encoder

We represent the dFC matrix **C**(*t_k_*) at each window *t_k_* as a weighted graph with *N* nodes corresponding to brain regions, in which edge (*i, j*) carries the correlation *C_ij_*(*t_k_*) and is retained when |*C_ij_*(*t_k_*)| ≥ 0.2. Each node is assigned a learnable 128-dimensional region embedding shared across subjects and windows, so that window-specific information enters the encoder only through the edge weights. A GATv2 graph attention network [24] with *L*_GNN_ = 4 layers encodes each graph through attention-weighted aggregation of neighboring node representations, with attention coefficients that depend jointly on the source and target node representations and the connecting edge weight, allowing the relative importance of neighboring regions to vary with the connectivity state at each window. The first three layers use four attention heads of width 128 with concatenated outputs, and the final layer uses a single head with output dimension *d_z_* = 256. Each layer is followed by layer normalization, with ELU activation and dropout after all but the final layer. Node representations from the final layer are mean-pooled across all *N* regions to yield a graph-level embedding at each window,

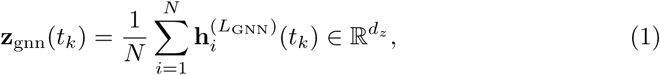

yielding the sequence 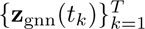. The time-averaged representation is obtained by averaging this sequence over windows and passing the result through a two-layer projection head, giving **z̄**_gnn_ ∈ ℝ^128^. The neural ODE is initialized from a single latent state **z**_0_ = Z0Enc(**z**_gnn_(*t*_1_)) ∈ ℝ*^d_z_^*, where Z0Enc is a three-layer MLP with hidden width 256 that maps the first window’s embedding into the ODE latent space. This anchors each trajectory to the observed initial connectivity state, while its subsequent evolution is governed by the learned dynamics.

#### 2.3.2 Neural ODE Dynamical System

Starting from the initial latent state **z**_0_, we model latent evolution using the autonomous ODE

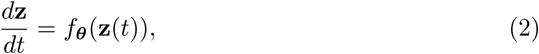

where 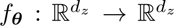 is a three-layer MLP with hidden width 512 and tanh activations. These activations yield a smooth, bounded vector field that supports stable adaptive integration. The latent trajectory is obtained by integrating the ODE,

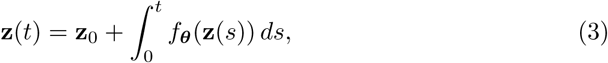

and evaluating the solution at *T* equally spaced points *t*_1_*,…, t_T_* on the normalized interval [0, 1] to obtain **Z**_traj_ ∈ ℝ*^T ×d_z_^*. We use the Dormand–Prince adaptive Runge–Kutta solver (dopri5) in torchdiffeq [22] with relative and absolute tolerances of 10^−3^ and 10^−4^, respectively. Gradients are computed by backpropagating directly through the solver operations rather than using the adjoint method, thereby differentiating the computed numerical trajectory. The vector field *f****_θ_*** is time-invariant and shared across all conditions and individuals within a dataset. Consequently, condition-specific and individual-specific behavior is encoded entirely in the initial state **z**_0_, while all trajectories evolve under the same learned dynamics.

#### 2.3.3 Trajectory Feature Extractor

The latent trajectory 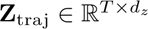 is summarized into a fixed-dimensional representation 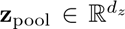 through two parallel streams. The *scalar stream* computes eight geometric descriptors, 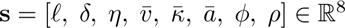, where 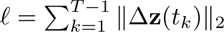 is the path length, 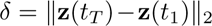 the net displacement, *η* = *ℓ/*(*δ* +*ε*) the tortuosity with 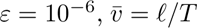 the mean speed, 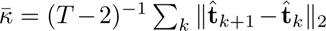 the mean curvature with unit tangent 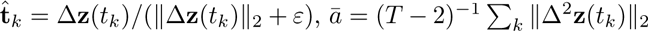 the mean acceleration, *ϕ* = max*_k_* ||**z**(*t_k_*) − **z̄**|| _2_ the peak excursion from the trajectory mean, and *ρ* the temporal speed asymmetry, defined as the mean step size in the first half of the trajectory minus that in the second half. Together, these describe the extent, efficiency, directional change, and temporal profile of state-space traversal, and are projected to ℝ*^d_z_^* by a linear layer. The *state stream* summarizes how the latent state changes across the trajectory by dividing it into three equal segments. The mean latent states of the early, middle, and late segments, **z**_early_, **z**_mid_, and **z**_late_, are concatenated with the per-dimension temporal variance **z**_var_ = Var*_k_*(**z**(*t_k_*)) and passed through a three-layer MLP with hidden width 512 and tanh activations, giving **z**_phase_ = MLP([**z**_early_ || **z**_mid_ || **z**_late_ || **z**_var_]) ∈ ℝ*^d_z_^*. The two streams are combined additively to form the trajectory representation **z**_pool_ = Linear(8*, d_z_*)(**s**) + **z**_phase_ ∈ ℝ*^d_z_^*.

#### 2.3.4 Classifier Head

The trajectory representation **z**_pool_ and the time-averaged representation **z̄**_gnn_ are concatenated and passed through a three-layer MLP classifier with hidden width 512, tanh activations, dropout *p* = 0.5, and a softmax output over the *M* conditions, and the predicted condition is the most probable output of

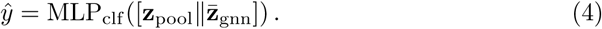

This design enables us to assess the contribution of each representation at inference by setting the other to zero, giving *ŷ*_traj_ = MLP_clf_ ([**z**_pool_ || **0**]) and *ŷ*_gnn_ = MLP_clf_ ([**0 || z̄**_gnn_]), which reflect the contribution of the trajectory and time-averaged representations, respectively. The difference between ablated and combined accuracy across conditions quantifies these contributions, providing the basis for testing the information migration principle. Full model hyperparameters and training details are reported in Supplementary A1.

#### 2.3.5 Loss Function

The model is trained by minimizing a composite objective that combines classification, contrastive, fitting, and alignment losses,

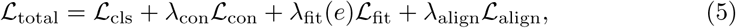

where *e* denotes the training epoch. The classification term *L*_cls_ is cross-entropy with label smoothing. The contrastive loss *L*_con_ is a supervised NT-Xent loss [32] applied to **z̄**_gnn_ with temperature *τ*_con_ = 0.1, encouraging same-condition samples to cluster in the time-averaged embedding space. The system identification loss *L*_fit_ is the mean squared error between ODE trajectory states and stop-gradient GNN embeddings, averaged over samples, windows, and latent dimensions,

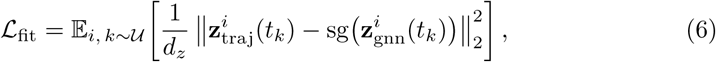

where sg(·) is the stop-gradient operator, so that the GNN embeddings act as a fixed target for the trajectory. This loss anchors the trajectory to the observed connectivity dynamics, and its weight is annealed from an initial value 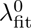 so that this constraint relaxes over training,

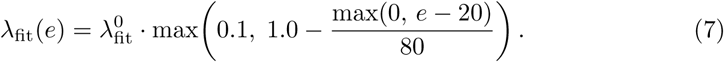

The cross-modal alignment loss *L*_align_ is a supervised NT-Xent loss between **z**_pool_ and **z̄**_gnn_, with the latter zero-padded to *d_z_* after L2 normalization. Same-condition pairs from different samples are treated as positives across the two representations, encouraging them to occupy consistent regions of a shared normalized embedding space.

### 2.4 Dynamical Metrics

We characterized the learned dynamics with three metrics, the ergodicity gap, the temporal variance ratio, and the participation ratio. The ergodicity gap, 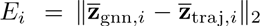 with 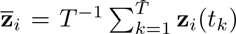, is the Euclidean distance between the mean of the observed embedding sequence of sample *i* and the mean of a trajectory generated by the learned dynamics, and thus reflects how strongly the observed temporal structure constrains those dynamics. A smaller gap indicates that the trajectory closely tracks the embeddings, as expected when the fitting loss constrains a trajectory through a structured sequence of distinct configurations, whereas a larger gap indicates weaker coupling, as expected when the embeddings remain near a single configuration and the trajectory is only weakly constrained. Smaller gaps therefore correspond to less ergodic, higher-complexity dynamics and larger gaps to more ergodic-like, lower-complexity dynamics, and we use the ergodicity gap as our measure of chronnectome complexity. It is a model-derived proxy for ergodic breakdown rather than a formal test of ergodicity. The temporal variance ratio, TVR*_i_* = Var*_t_*(**z**_traj*,i*_)*/*(Var*_t_*(**z**_gnn*,i*_) + *ε*), with variance computed over time points and averaged over latent dimensions, measures whether the learned dynamics amplify (TVR *>* 1) or suppress (TVR *<* 1) temporal variance relative to the embeddings. The dimensionality of the latent space occupied by each condition is quantified by the participation ratio [33], 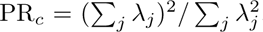, where *λ_j_* are the eigenvalues of the covariance of the trajectory states {**z**_traj*,i*_(*t_k_*)} pooled across all time points and all samples *i* of condition *c*, so that higher values indicate trajectories that together spread across more latent dimensions. Participation ratio thus measures the breadth of the dynamics (dynamical richness), a property distinct from their sequential structure and therefore from chronnectome complexity. Differences in participation ratio between groups were assessed with 90% confidence intervals from bootstrap resampling of samples. Full definitions, statistical procedures, and additional metrics are provided in Supplementary A5.

## 3 Results

Results are reported across four experimental configurations. Experiment (A) used HCP-YA with the cortical atlas, Experiment (B) used HCP-YA with the cortico-subcortical atlas, Experiment (C) used HCP-Aging with the cortical atlas, and Experiment (D) used HCP-Aging with the cortico-subcortical atlas. All evaluations used held-out test sets partitioned at the subject level. We first validate the model’s classification performance, then use inference-time ablation to examine where condition-discriminative information resides and the ergodicity gap to relate this organization to chronnectome complexity, and finally ask how it depends on subcortical coverage and on age. Dynamical metrics are defined in Methods, with full definitions and statistical procedures in Supplementary A5 and A8.

### 3.1 Classification Performance

The model achieved high classification accuracy in all four experiments, evaluated on held-out test sets partitioned at the subject level (Table 2). In HCP-YA, accuracy reached 97.40% for the cortical atlas and 94.20% for the cortico-subcortical atlas, against a chance level of 12.5% for eight classes. In HCP-Aging, accuracy was 92.22% for the cortical atlas and 92.29% for the cortico-subcortical atlas, against a chance level of 25% for four classes. Per-condition precision, recall, and F1 scores for the full model, together with confusion matrices, are provided in Supplementary A7.

**Table 2.** Classification performance across experiments. The cortico-subcortical atlas comprises the 100 Schaefer cortical regions and the 16 subcortical regions of Scale I of the Melbourne Subcortex Atlas. Parentheses show 95% confidence intervals.

| Experiment | Acc (%) | Cohen's *κ* | MCC | AUC | F1 | Precision | Recall |
|---|---|---|---|---|---|---|---|
| (A) HCP-YA<br>Cortical | 97.40<br>(96.6–98.2) | 0.970<br>(0.961–0.979) | 0.970<br>(0.961–0.979) | 0.999 | 0.974 | 0.974 | 0.974 |
| (B) HCP-YA<br>Cortico-subcortical | 94.20<br>(93.0–95.3) | 0.934<br>(0.920–0.946) | 0.935<br>(0.922–0.947) | 0.998 | 0.942 | 0.948 | 0.942 |
| (C) HCP-Aging<br>Cortical | 92.22<br>(90.4–94.0) | 0.896<br>(0.872–0.920) | 0.897<br>(0.873–0.920) | 0.991 | 0.922 | 0.923 | 0.922 |
| (D) HCP-Aging<br>Cortico-subcortical | 92.29<br>(90.5–93.9) | 0.897<br>(0.874–0.918) | 0.900<br>(0.879–0.920) | 0.992 | 0.924 | 0.933 | 0.923 |

For context, we compared the model with published methods evaluated on the same HCP-YA cohort (Table 3). Although our evaluation setting was more challenging, using a coarser parcellation of 100 rather than 400 regions and a model trained on eight classes including resting state, the model performed close to dynamic methods and above static ones, reaching 97.03% accuracy on the seven task conditions without retraining. These results establish the model as a reliable classifier of resting-state and task conditions and provide a validated basis for the ablation analyses that follow.

**Table 3.** Comparison with published methods on HCP-YA task classification. Published results are from Kim et al. [19], evaluated on seven task classes without resting state (mean *±* s.d. across cross-validation folds), while our model was evaluated on a held-out test set (bootstrap s.d.).

| Method | Type of Input | ROIs | Accuracy (%) |
|---|---|---|---|
| STAGIN-SERO [19] | Dynamic FC | 400 | 99.19 ± 0.20 |
| STAGIN-GARO [19] | Dynamic FC | 400 | 99.02 ± 0.17 |
| ST-GCN [18] | Dynamic FC | 400 | 98.92 ± 0.27 |
| BAnD [34] | BOLD | — | 95.10 ± 0.62 |
| GIN [35] | Static FC | 400 | 93.87 ± 0.66 |
| ChebGCN [36] | Static FC | 400 | 73.06 ± 0.68 |
| GraphSAGE [37] | Static FC | 400 | 54.52 ± 0.97 |
| GCN [38] | Static FC | 400 | 45.07 ± 1.63 |
| Ours (7 task classes) | Dynamic FC | 100 | 97.03 ± 0.44 |
| Ours (8 classes) | Dynamic FC | 100 | 97.40 ± 0.41 |

### 3.2 Information Migration Across Conditions

The information migration principle predicts that trajectory geometry should carry more condition-discriminative information under higher-complexity conditions, with time-averaged connectivity becoming an insufficient descriptor of brain state as chronnectome complexity increases. To test this, we performed complementary ablations at inference time, setting either the time-averaged or the trajectory representation to zero while preserving the other, and measured the resulting classification accuracy for each condition. The significance of trajectory-only accuracy was assessed by label permutation, as described in Supplementary A3, and chronnectome complexity was quantified by the ergodicity gap.

Trajectory-only accuracy in Experiment (A) spanned a wide range across conditions, from 93.4% for Working Memory to 0.0% for Relational Reasoning, which was indistinguishable from the label-permutation null (Table 4 and Fig. 2a). Resting-state trajectories were also classified with high accuracy (89.3%), although rest is among the most ergodic conditions, likely reflecting their distinct location in the latent space and high dynamical richness rather than a structured sequence of configurations. In Experiment (B), the extremes of this range were preserved, with Working Memory remaining high (88.3%) and Relational Reasoning at 0.0%. Relational Reasoning remained at or near 0.0% across all loss configurations tested (Supplementary A7), because its trajectories closely resemble those of Working Memory, consistent with shared frontoparietal control demands. These results reveal a division of labor between the two representations. Under the non-ergodic Working Memory condition, the trajectory representation captured condition identity, as reflected in its high trajectory-only classification accuracy. For conditions whose trajectories were not distinctive, and whose samples were therefore assigned to Working Memory under trajectory-only classification, the time-averaged representation supplied the information the full model used to classify them. This was clearest for Relational Reasoning, the most ergodic condition, which was classified almost perfectly by the full model (99.5%) but not at all from its trajectory alone, consistent with its condition-discriminative information residing in time-averaged connectivity.

**Fig. 2.**
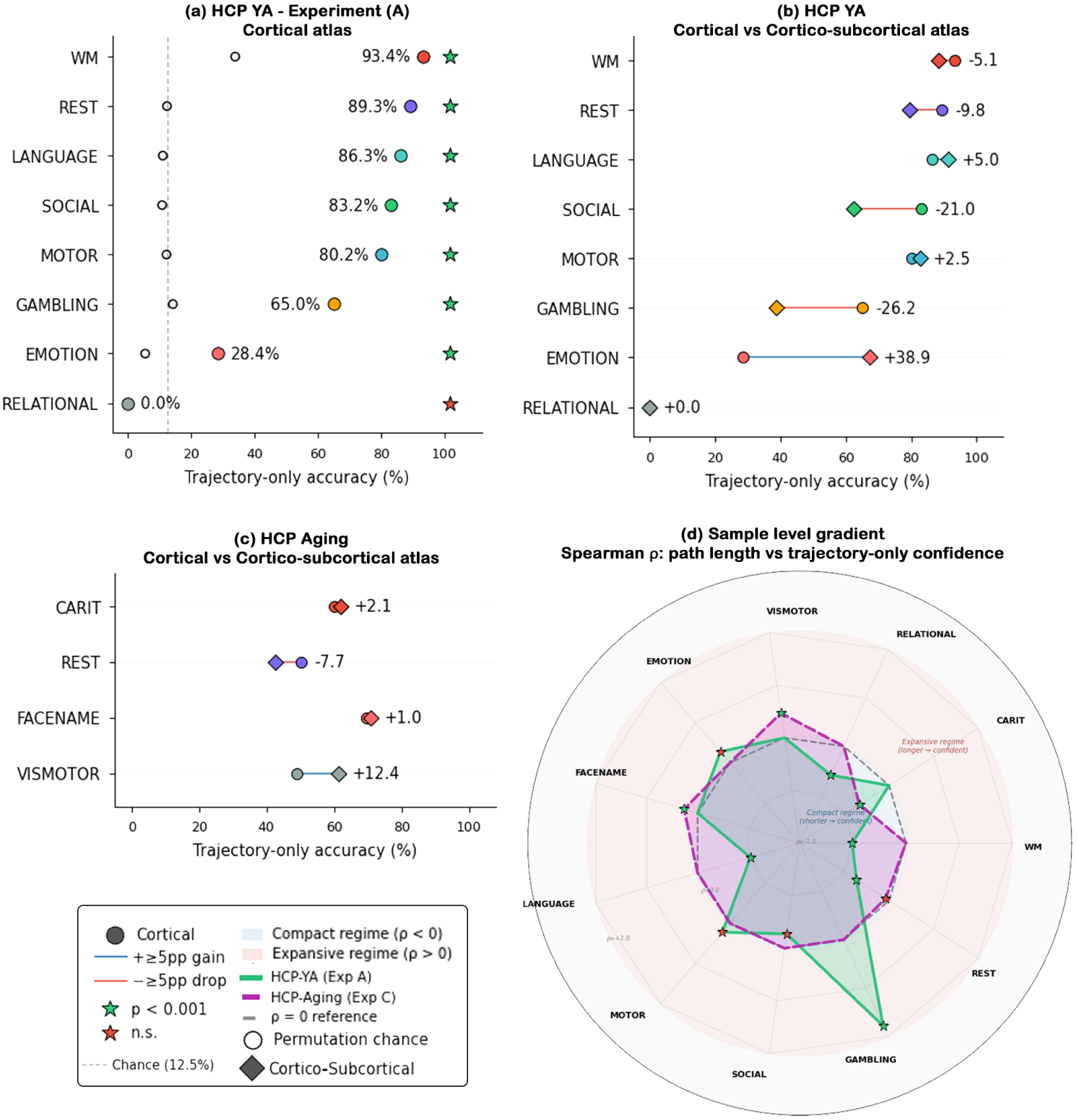
Trajectory-only accuracy varies across conditions, cohorts, and atlas configurations. (**a**) Per-condition trajectory-only accuracy for the HCP-YA cortical atlas (Experiment A). The dashed line indicates the chance baseline. (**b**,**c**) Paired comparison between cortical and cortico-subcortical atlas configurations for HCP-YA and HCP-Aging, showing condition-specific, atlas-dependent changes. (**d**) Spearman *ρ* between trajectory path length and trajectory-only classification confidence for each condition. Blue inner zone, compact regime (*ρ <* 0); red outer zone, expansive regime (*ρ >* 0). Green and red stars indicate significant and non-significant effects, respectively.

**Table 4.** Per-condition classification accuracy (%) of the full model and of trajectory-only classification, obtained by setting the time-averaged representation to zero at inference. Columns correspond to Experiments (A) and (B) for HCP-YA and (C) and (D) for HCP-Aging.

|  | **HCP-YA** |  |  |  |  | **HCP-Aging** |  |  |  |
|---|---|---|---|---|---|---|---|---|---|
|  | **Cortical** |  | **Cortico-subcortical** |  |  | **Cortical** |  | **Cortico-subcortical** |  |
| **Condition** | Full | Traj. | Full | Traj. | **Condition** | Full | Traj. | Full | Traj. |
| WM | 93.9 | 93.4 | 81.1 | 88.3<sup>*b*</sup> | CARIT | 84.4 | 60.0 | 82.5 | 62.1 |
| REST | 100.0 | 89.3 | 100.0 | 79.6 | REST | 95.6 | 50.2 | 98.8 | 42.5 |
| LANGUAGE | 99.5 | 86.3 | 99.0 | 91.3 | FACENAME | 93.8 | 69.8 | 90.8 | 70.8 |
| SOCIAL | 99.5 | 83.2 | 98.0 | 62.2 | VISMOTOR | 95.1 | 48.9 | 97.1 | 61.3 |
| MOTOR | 100.0 | 80.2 | 99.0 | 82.7 |  |  |  |  |  |
| GAMBLING | 90.4 | 65.0 | 80.1 | 38.8 |  |  |  |  |  |
| EMOTION | 96.4 | 28.4 | 99.0 | 67.3 |  |  |  |  |  |
| RELATIONAL | 99.5 | 0.0<sup>*a*</sup> | 97.4 | 0.0<sup>*a*</sup> |  |  |  |  |  |

<sup>*a*</sup>Indistinguishable from the label-permutation null. <sup>*b*</sup>Exceeds full-model accuracy because samples from other conditions are also assigned to Working Memory under trajectory-only classification.

Two training ablations on the HCP-YA cortical atlas tested how the loss terms shape these results. A model trained without the fitting loss (*λ*_fit_ = 0), with all other objectives unchanged, preserved the overall pattern, with trajectory-only accuracy changing by less than 8 percentage points for most conditions. The exception was Gambling, whose accuracy fell from 65.0% to 31.5%, indicating that the fitting loss specifically shapes trajectory geometry for this condition, which also showed the strongest within-condition relationship between trajectory path length and trajectory-only classification confidence. The cross-modal alignment loss is integral to the design, organizing the trajectory and time-averaged representations by condition in a shared space so that the classifier draws on both, and without it (*λ*_align_ = 0) overall accuracy was largely preserved (95.75%) but the classifier did not fully capture trajectory dynamics (Supplementary A.2). A linear probe trained directly on the trajectory representation of this model nevertheless reached 70.2% accuracy, compared with 77.8% with alignment, with a similar pattern across conditions. The trajectory representation therefore carries condition-discriminative information without alignment, and the alignment loss mainly helps the model use it.

In HCP-Aging (Experiments C and D), trajectory-only accuracy was lower overall than in HCP-YA despite a higher chance level, and the time-averaged representation carried most of the condition-discriminative information for every condition (Table 4), although the contrast between higher- and lower-complexity conditions was preserved in the cortical atlas, with CARIT exceeding Visuomotor Tracking (60.0% against 48.9%). Across both cohorts, the relationship between trajectory path length and trajectory-only classification confidence within each condition revealed two regimes (Fig. 2d). Shorter trajectories were associated with higher confidence for Working Memory, Relational Reasoning, and Rest in both atlases, and for Social and CARIT in one (*ρ* from −0.17 to −0.71), whereas longer trajectories were associated with higher confidence for Gambling and Face-Name Memory in both atlases and for Emotion with subcortical inclusion (*ρ* from +0.13 to +0.89). Language and Visuomotor Tracking changed regime with the atlas, indicating that for some conditions, which trajectories carry condition identity depends on whether subcortical regions are included.

### 3.3 Subcortical Observability Boundary

Extending connectivity to include subcortical regions in Experiments (B) and (D) produced condition-specific changes in trajectory contributions (Fig. 2b,c), identifying an anatomical observability boundary for conditions whose dynamics depend on subcortical circuits. In Experiment (B), Emotion trajectory-only accuracy increased from 28.4% to 67.3%, consistent with amygdala dynamics lying outside the observability window of cortical parcellations, and the relationship between its trajectory path length and classification confidence shifted from non-significant (*ρ* = +0.138) to strongly positive (*ρ* = +0.629, *p <* 0.001). Gambling showed the opposite effect, with higher dynamical richness (participation ratio 7.54 to 9.13) but lower trajectory-only accuracy (65.0% to 38.8%), suggesting that striatal dynamics broaden the explored state space while adding within-condition variability that reduces condition separability. For Working Memory, subcortical inclusion reduced the information available from time-averaged connectivity, which identified it in 88.3% of samples in Experiment (A) but only 54.1% in Experiment (B), while trajectory-only accuracy remained high in both (93.4% and 88.3%). These condition-specific effects indicate that the observability boundary depends on the circuits involved rather than reflecting a uniform effect of atlas expansion. In HCP-Aging, Visuomotor Tracking gained 12.4 percentage points with subcortical inclusion (48.9% to 61.3%), whereas Face-Name Memory changed little (69.8% to 70.8%).

### 3.4 Chronnectome Complexity and Ergodicity Gap

The ergodicity gap compares the mean of the embedding sequence with the mean of the trajectory generated by the learned dynamics, providing a model-derived index of ergodic breakdown that we use to quantify chronnectome complexity (Supplementary A5). In Experiment (A), Working Memory had the smallest ergodicity gap (9.00 ± 0.79) and Relational Reasoning the largest (9.89 ± 0.83), identifying them as the highest- and lowest-complexity conditions (Mann–Whitney *U* = 8,473, *p* = 3.99×10^−22^, *d* = −1.09; Fig. 3a). The full ordering from smallest to largest gap was Working Memory (9.00), Gambling (9.29), Language (9.44), Emotion (9.53), Motor (9.73), Social (9.83), Rest (9.86), and Relational Reasoning (9.89). The two conditions at the ends of this ordering were also those at the ends of the trajectory-only range, with Working Memory identified by its trajectory and Relational Reasoning by time-averaged connectivity, whereas Rest combined a large gap with high trajectory-only accuracy, as discussed above.

**Fig. 3.**
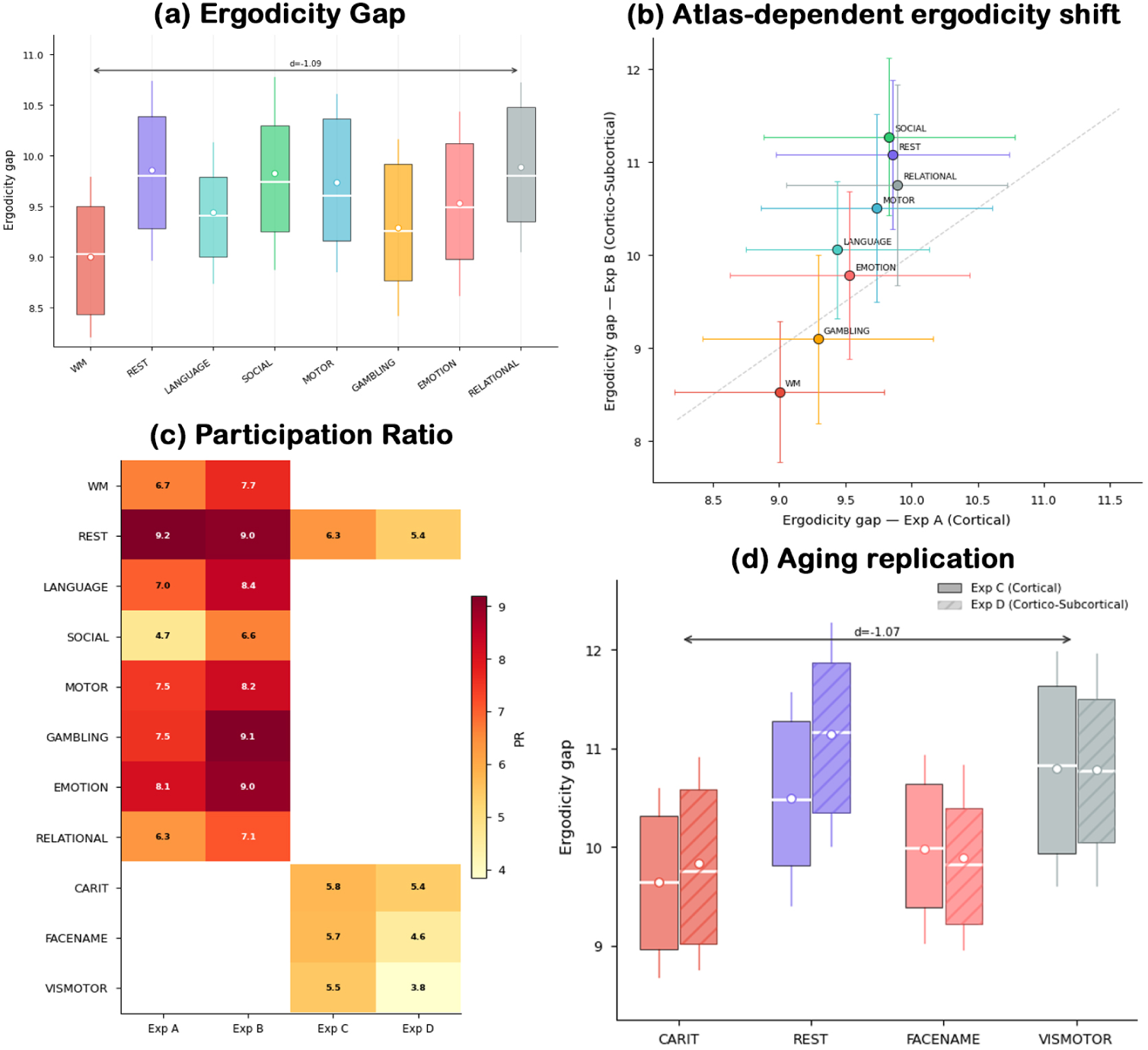
Ergodicity gap and dynamical richness across experiments. (**a**) Ergodicity gap for each condition in Experiment (A), shown as box plots (interquartile range with median line, and whiskers showing mean *±* s.d.). Smaller gaps indicate tighter coupling between the learned trajectory and the embeddings and higher chronnectome complexity. (**b**) Mean ergodicity gap for each condition in Experiment (A) against Experiment (B), with error bars showing *±* s.d. and the dashed identity line for reference. Because the two experiments were trained separately, the relative ordering of conditions within each experiment is informative, whereas absolute values are not directly comparable. (**c**) Dynamical richness (participation ratio) for each condition across the four experiments, each computed within its own model. (**d**) Ergodicity gap distributions for HCP-Aging in the cortical (Experiment (C), solid) and cortico-subcortical (Experiment (D), hatched) atlas configurations.

**Fig. 4.**
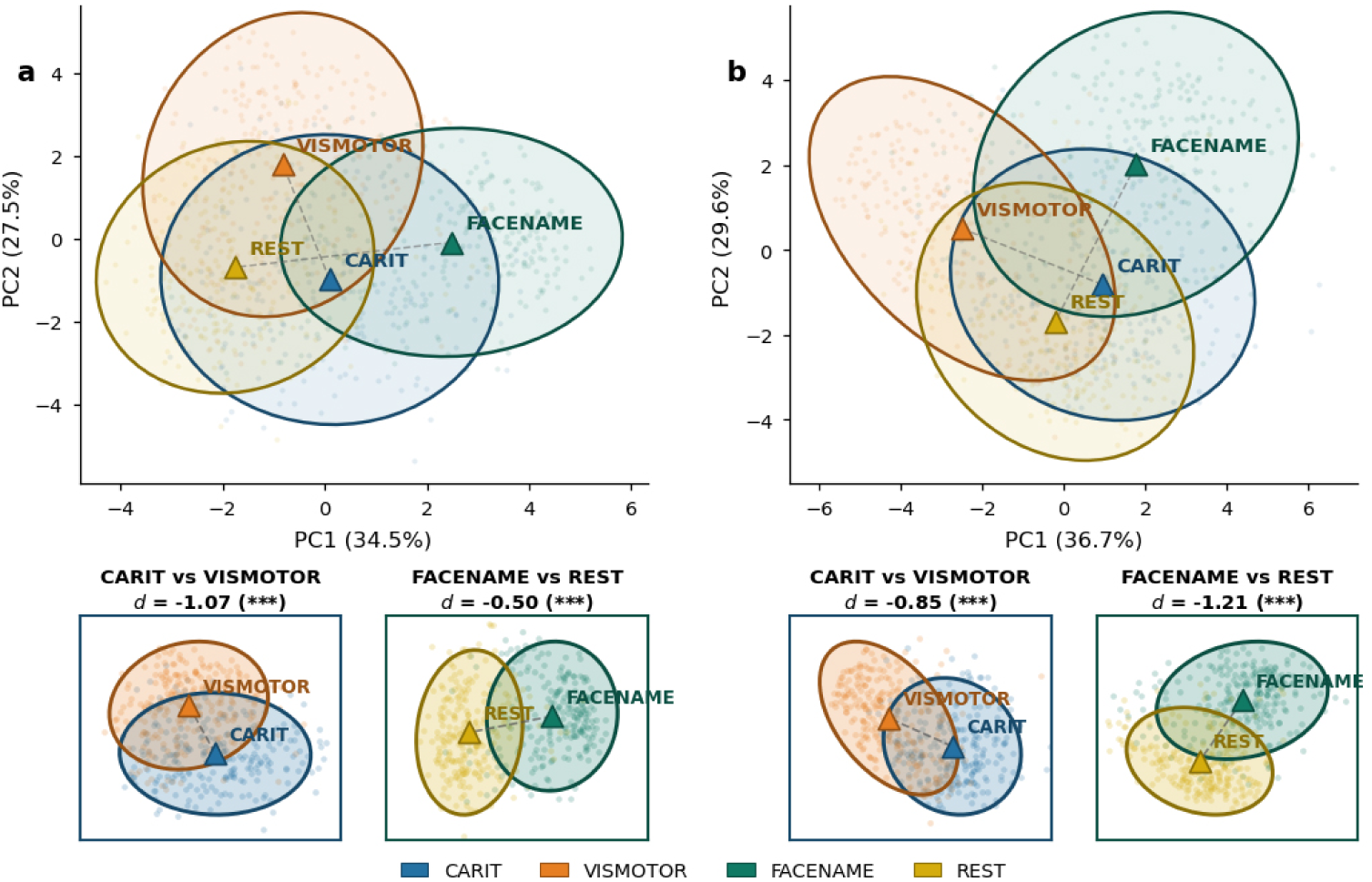
Latent trajectory organization within the HCP-Aging cohort. (**a**) Cortical atlas, Experiment (C). (**b**) Cortico-subcortical atlas, Experiment (D). In each column, the upper panel shows the first two principal components of each sample’s time-averaged trajectory for the four HCP-Aging conditions, with ellipses covering 2 s.d. of the samples, triangles marking condition centroids, and dashed lines connecting the centroids of each compared pair. The lower panels enlarge the comparisons between CARIT and Visuomotor Tracking (left) and between Face-Name Memory and Rest (right). Values of *d* give Cohen’s *d* for the difference in ergodicity gap between the two conditions, with negative values indicating a smaller gap, and therefore higher chronnectome complexity (*^∗∗∗^p <* 0.001, Mann–Whitney test). Principal components were computed separately for each atlas configuration, so positions are comparable within but not across columns.

In Experiment (B), the contrast between Working Memory and Relational Reasoning more than doubled in effect size (*d* = −2.39, *p* = 9.64 × 10^−53^; Fig. 3b), with gaps of 8.53 and 10.75, respectively. We compare effect sizes rather than absolute gap values across atlases, since Experiments (A) and (B) were trained separately. In HCP-Aging, CARIT showed a smaller gap than Visuomotor Tracking in both the cortical (*d* = −1.07, *p* = 8.58 × 10^−24^) and cortico-subcortical atlases (*d* = −0.85, *p* = 6.12 × 10^−18^; Fig. 3d), replicating the contrast between higher- and lower-complexity conditions in an independent cohort. In Experiment (D), the temporal variance ratio was perfectly inversely ranked with trajectory-only accuracy across the four conditions (*ρ* = −1.0), consistent with near-stationary conditions, whose embeddings vary little over time, carrying their condition-discriminative information in time-averaged connectivity, although four conditions provide limited statistical power.

### 3.5 Dynamical Richness and Energy Proxy

Dynamical richness, measured by participation ratio, differed across conditions within each cohort (Fig. 3c). In HCP-YA, Rest showed the highest dynamical richness in the cortical atlas (9.20), indicating that its trajectories together spread across the most latent dimensions despite low chronnectome complexity, whereas Social showed the lowest (4.71). Subcortical inclusion, compared across separately trained models, affected the two cohorts in opposite directions. In HCP-YA, participation ratio increased for every condition except Rest, most strongly for Social (4.71 to 6.64) and Gambling (7.54 to 9.13), whereas in HCP-Aging it decreased for all four conditions, most strongly for Visuomotor Tracking (5.47 to 3.84), suggesting that subcortical circuits broaden the explored state space in young adults but operate within a narrower dynamical range in older adults. As an energy proxy, we used trajectory path length, under the assumption that longer traversal of the latent space reflects greater dynamical cost (Fig. 5c). In HCP-YA, path length was negatively correlated with mean absolute functional connectivity amplitude across all samples (Experiment (A), *r* = −0.182, *p* = 3.33×10^−13^; Experiment (B), *r* = −0.188, *p* = 5.83×10^−14^). In HCP-Aging, this relationship was not significant across all samples (*r* = +0.002, *p* = 0.947 in Experiment (C); *r* = +0.051, *p* = 0.116 in Experiment (D)), whereas Face-Name Memory (*r* = +0.151, *p* = 0.023) and Visuomotor Tracking (*r* = +0.179, *p* = 0.007) showed positive correlations in Experiment (C), indicating that longer trajectories in aging were associated with stronger mean connectivity for these conditions.

**Fig. 5.**
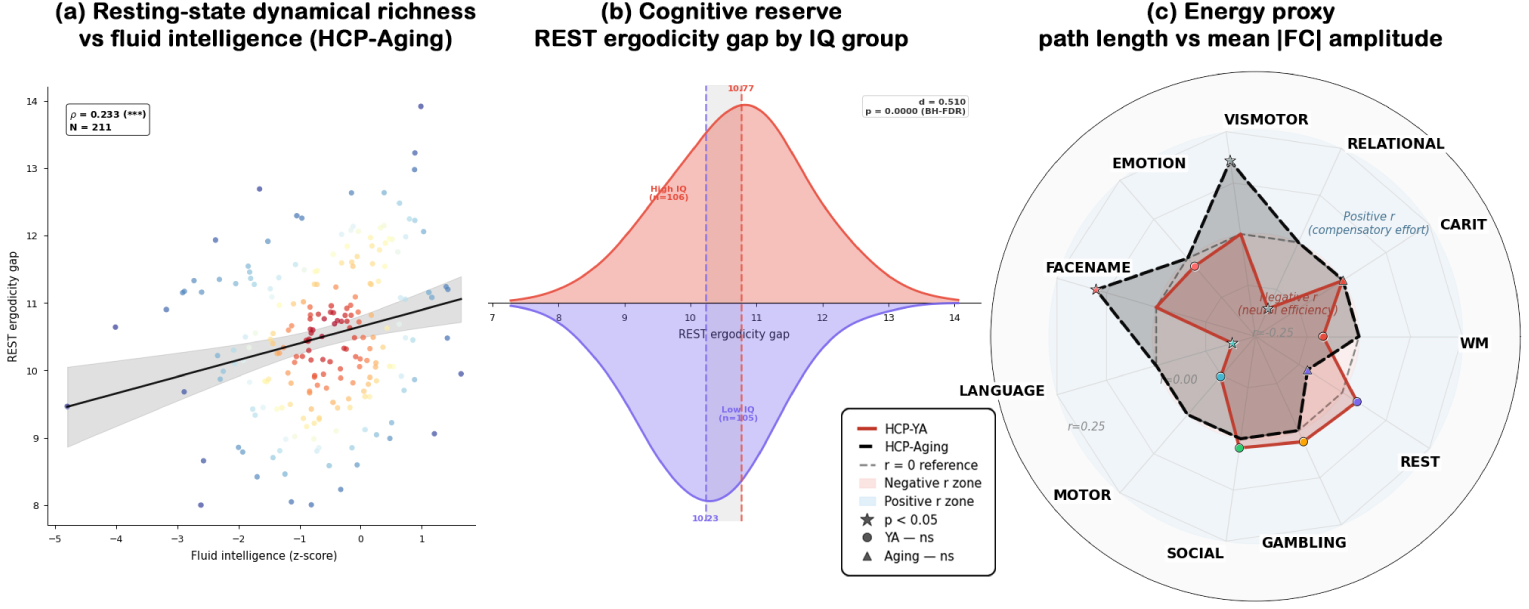
Resting-state ergodicity, cognitive reserve, and the energy proxy. (**a**) Correlation between the resting-state ergodicity gap and fluid intelligence in HCP-Aging (Experiment (C), *ρ* = 0.233, *p* = 0.0006, *N* = 211), with regression line and 95% confidence band. Points are colored by local density. (**b**) Resting-state ergodicity gap by median-split fluid intelligence group in HCP-Aging (Experiment (C)), shown as a split density plot with high-IQ (red) and low-IQ (blue) distributions mirrored about the horizontal axis. The dashed vertical line marks the grand mean, and the effect size and BH-FDR corrected *q*-value are annotated. (**c**) Pearson *r* between trajectory path length and mean absolute functional connectivity amplitude for each condition, for HCP-YA (red polygon, Experiment (A)) and HCP-Aging (blue dashed polygon, Experiment (C)). The dashed ring marks *r* = 0, with the inner zone indicating negative and the outer zone positive correlations.

### 3.6 Information Migration in Aging

We assessed aging by testing whether the contrasts observed in young adults were reproduced within the aging cohort, which includes its own higher- and lower-complexity conditions, so that conditions are compared under the same learned dynamics. The distinction between higher- and lower-complexity conditions was preserved, with CARIT showing the smallest and Visuomotor Tracking the largest ergodicity gap among HCP-Aging task conditions in both atlases (Fig. 3d), paralleling the contrast between Working Memory and Relational Reasoning in HCP-YA (Fig. 3a), and the two conditions also occupied distinct regions of each model’s latent space (Fig. 4a,b). Face-Name Memory likewise showed a smaller gap than Rest (*d* = −0.50 and −1.21, both *p <* 0.001). CARIT also showed higher trajectory-only accuracy than Visuomotor Tracking in the cortical atlas (60.0% against 48.9%), consistent with the information migration principle, and remained marginally higher with subcortical inclusion (62.1% against 61.3%), where Visuomotor Tracking gained 12.4 percentage points, consistent with thalamocortical contributions to visuomotor dynamics that are not observable in the cortical atlas. The time-averaged representation nonetheless carried most of the condition-discriminative information for all HCP-Aging conditions. Overall, dynamical richness was lower in HCP-Aging across all matched conditions and both atlases, although this comparison is based on separately trained models and therefore cannot be attributed to aging alone.

For resting state, the ergodicity gap decreased with age within HCP-Aging (*ρ* = −0.158, *p* = 0.022), indicating that resting-state dynamics departed from their near-ergodic regime in older participants, a change that reflects a loss of stationary organization rather than the structured sequencing that characterizes higher-complexity tasks. At the same time, the resting-state ergodicity gap was positively correlated with fluid intelligence (*ρ* = 0.233, *p* = 0.0006, *N* = 211; Fig. 5a), and this association remained significant after controlling for age (partial *ρ* = 0.178, *p* = 0.0096), so that older adults with higher fluid intelligence had more ergodic resting-state dynamics independent of age. A median-split comparison, unadjusted for age, was consistent with this result (high-IQ mean = 10.77, low-IQ mean = 10.23, *d* = 0.51, *p*_perm_ *<* 0.001, *q <* 0.001, BH-FDR across 12 tests; Fig. 5b). The effect was absent in Experiment (D) (*d* = 0.09, *p* = 0.506, *q* = 0.905), indicating that it was specific to the cortical atlas. Cross-validated prediction of fluid intelligence from trajectory features and group comparisons across all conditions and cohorts are reported in Supplementary A6.

## 4 Discussion

The principal finding of this work is that chronnectome complexity is associated with where condition-discriminative information resides in brain dynamics, supporting the information migration principle. Under Working Memory, the least ergodic condition, the trajectory carried condition identity, whereas under Relational Reasoning, the most ergodic condition, condition identity resided almost entirely in time-averaged connectivity, so that the two representations derived from a single learned dynamical system divided the information between them. The results further indicate that chronnectome complexity is distinct from cognitive demand and conventional task difficulty. Despite its cognitive demands, Relational Reasoning exhibited near-ergodic behavior, with trajectories that closely resembled those of Working Memory and carried little condition-specific information. This pattern is consistent with sustained frontoparietal engagement within a relatively stable configuration. In contrast, the n-back Working Memory task requires continuous encoding, comparison, and updating of items, which may impose a more structured sequence of brain configurations, breaking ergodicity. Training ablations further showed that the trajectory representation carries condition-discriminative information even without the alignment loss, and that this loss mainly determines whether the classifier draws on that information.

Resting state is often described as the most complex brain state, in the sense of the entropy or breadth of its dynamical repertoire [39, 40]. Our results agree with this view, as Rest showed among the highest dynamical richness of all conditions, yet it showed low chronnectome complexity, indicating that breadth of exploration and sequential organization are distinct properties of brain dynamics. Our data show this dissociation directly, as Working Memory and Relational Reasoning had similar dynamical richness but lay at opposite ends of the ergodicity gap, whereas Rest combined the highest richness with one of the largest ergodicity gaps. The breadth of resting dynamics is consistent with evidence that spontaneous activity explores a broad repertoire of network configurations in the vicinity of multistable attractors [12], and that task engagement reduces the variability of large-scale activity relative to rest [41]. This stabilizing effect of task engagement concerns the breadth of activity, reflected in dynamical richness, rather than its sequential organization, reflected in the ergodicity gap, so that tasks may constrain activity while still sequencing it through distinct regimes. At the same time, resting dynamics were near-ergodic, consistent both with evidence that resting-state connectivity is largely described by a single stable correlation structure [42] and with evidence that it alternates among a set of recurring connectivity states [10]. Even the sequential structure reported in resting-state dynamics [43] is compatible with near-ergodicity, since sequentially correlated exploration that repeatedly revisits the same connectivity configurations still yields time averages that represent the dynamics, whereas ergodic breakdown arises when activity diverges from a common attractor repertoire, so that the time average conflates the distinct configurations it visits. This near-ergodic regime also appears functionally relevant, as its preservation in older adults was associated with higher fluid intelligence.

Including subcortical regions changed trajectory contributions in condition-specific ways, indicating that cortical parcellations miss part of the dynamics of conditions that depend on subcortical circuits. The increase for Emotion is consistent with amygdala contributions to affective processing [44], and the reduced separability of Gambling despite its broader spread across the latent space is consistent with striatal reward dynamics varying across samples, making Gambling trajectories more diverse and therefore harder to distinguish from other conditions [45]. For Working Memory, sub-cortical inclusion reduced the information available from time-averaged connectivity while leaving the condition identifiable from its trajectory. This pattern is consistent with the role of thalamo-prefrontal interactions in working memory [46], which unfold over the course of a task and would therefore be expressed in trajectory geometry rather than in temporal averages. In HCP-Aging, the increase in trajectory-only accuracy for Visuomotor Tracking is consistent with the engagement of the striatum and thalamus during visually guided tracking [47, 48], whereas the small change for Face-Name Memory suggests that the hippocampal dynamics engaged by associative encoding are already partly reflected in parahippocampal cortical regions, which are co-activated with the hippocampus during face-name encoding [49, 50], although the low signal-to-noise ratio of hippocampal fMRI signals may also limit their contribution. Trajectory length and trajectory-only classification confidence provide a complementary view of these conditions, suggesting a distinction between sustained and transient task responses. In Working Memory and Relational Reasoning, which involve predominantly sustained engagement, the most confidently classified samples followed relatively compact trajectories. In contrast, in Gambling, Face-Name Memory, and Emotion with subcortical regions included, the most confidently classified samples followed longer trajectories, in line with transient responses to reward outcomes, novel associations, and emotional faces. These events engage striatal, hippocampal, and amygdala circuits associated with reward processing, novelty, and emotional salience, respectively. This pattern is consistent with the distinction between sustained and transient components of task-evoked activity [51].

Within the older cohort, the distinction between higher- and lower-complexity conditions was preserved, with CARIT showing a smaller ergodicity gap and more trajectory-carried information than Visuomotor Tracking, indicating that the relationship between chronnectome complexity and the locus of condition-discriminative information extends into older age. The time-averaged representation nevertheless carried most of the condition-discriminative information for all HCP-Aging conditions, suggesting a weaker shift into trajectory geometry in older adults, although comparisons between cohorts involve separately trained models and cannot be attributed to aging alone. For resting state, the ergodicity gap decreased with age, indicating that resting-state dynamics depart from their near-ergodic regime in older participants, whereas higher fluid intelligence was associated with more ergodic resting-state dynamics, consistent with the view that cognitive reserve in aging is reflected in the maintenance of youthful patterns of brain function [52]. Finally, the energy proxy related trajectory path length negatively to mean connectivity amplitude in young adults, so that trajectories that traveled farther through the latent space were accompanied by weaker overall connectivity. This pattern suggests that an efficient brain moves between states by engaging specific regions at specific times rather than keeping all regions strongly coupled at once, consistent with accounts linking efficient neural computation to selective rather than global engagement [53]. In aging, this relationship was absent overall, whereas for Face-Name Memory and Visuomotor Tracking longer trajectories were accompanied by stronger overall connectivity. This pattern may reflect compensation [54], in which older adults recruit stronger large-scale coupling to support task performance, although the energy proxy remains on the assumption that longer trajectories reflect greater dynamical cost.

This work has some limitations, which also suggest directions for future research. The ergodicity gap is a model-derived proxy for ergodic breakdown rather than a formal test of ergodicity, and its interpretation depends on how strongly the fitting loss couples the trajectory to the observed embeddings, although the contrast between higher- and lower-complexity conditions it revealed, replicated across atlas configurations and in an independent cohort. It is common practice to assess the contribution of a representation by setting it to zero at inference, although this produces inputs outside the training distribution that can affect the result. In our analysis, this limitation was mitigated by label-permutation testing and by linear probes showing that condition-discriminative information is present in the trajectory representation itself. The two cohorts were modeled separately, by models trained on different numbers of conditions, so aging was assessed through comparisons within the aging cohort, and joint modeling of both cohorts would allow direct comparisons across the lifespan. Finally, the subcortical extension identifies circuit-level contributions without isolating specific nuclei, and higher-resolution subcortical parcellations, together with acquisitions optimized for subcortical signal, would sharpen these inferences.

Together, these findings support the information migration principle, under which the locus of condition-discriminative information shifts with chronnectome complexity from time-averaged connectivity to trajectory geometry. Under lower-complexity conditions, near-ergodic dynamics allow time averages to represent brain state faithfully, whereas under higher-complexity conditions, ergodic breakdown turns the time average into a ghost state and leaves the distinguishing information in the trajectory. Including subcortical regions modulated this organization in a condition-specific manner, and the organization was preserved in older adults. At rest, more ergodic dynamics were associated with higher fluid intelligence in aging, linking chronnectome complexity to the maintenance of cognitive function. These results suggest that analyses of brain dynamics should consider not only which connectivity states occur but also how their sequential structure determines where cognitive state information resides, opening several directions for future work, including extending chronnectome complexity to its spatial counterpart, modeling stochastic contributions to latent dynamics, and examining how ergodic breakdown is altered in development, neurodegeneration, and psychiatric conditions.

## Appendix A Supplementary Materials

### A.1 Architecture Hyperparameters and Training Details

All experiments used identical architecture hyperparameters except for the dataset-specific parameters listed in Table A1. The spatial encoder used 4 GATv2 layers with 4 attention heads, hidden dimension *d*_GNN_ = 128, dropout *p* = 0.2, and edge threshold *τ* = 0.2. The Z0 encoder used a three-layer MLP with hidden dimension 256. The neural ODE dynamics MLP used three layers with hidden dimension 512 and tanh activations. The trajectory feature extractor state stream used a three-layer MLP with hidden dimension 512. The classifier used a three-layer MLP with hidden dimension 512 and dropout *p* = 0.5. Total trainable parameters: 3,673,736.

**Table A1.** Experiment-specific configuration and training outcomes.

| **Exp.** | **ROIs** | **Classes** | *T*<sub>max</sub> | *λ*<sub>fit</sub> | *λ*<sub>align</sub> | **Best Epoch** | **Val Acc** |
|---|---|---|---|---|---|---|---|
| Exp A | 100 | 8 | 147 | 0.05 | 0.5 | 50 | 0.970 |
| Exp B | 116 | 8 | 147 | 0.05 | 0.5 | 38 | 0.966 |
| Exp C | 100 | 4 | 155 | 0.05 | 0.5 | 47 | 0.951 |
| Exp D | 116 | 4 | 155 | 0.05 | 0.5 | 47 | 0.931 |
| Exp A, no fit | 100 | 8 | 147 | 0.00 | 0.5 | 35 | 0.964 |
| Exp A, no align | 100 | 8 | 147 | 0.05 | 0.0 | 54 | 0.970 |

All other hyperparameters were identical across experiments. No fit and no align denote the training ablations without the fitting loss (*λ*<sub>fit</sub> = 0) and without the alignment loss (*λ*<sub>align</sub> = 0), respectively.

#### A.1.1 Training procedure

Models were trained for a maximum of 150 epochs with early stopping on validation loss. Optimization used AdamW with initial learning rate *η* = 2 × 10^−4^ and weight decay 10^−4^, with OneCycleLR scheduling stepped per batch. Training used Distributed Data Parallel across 8 NVIDIA A40 GPUs (46 GB each) with effective batch size 128 (16 per GPU × 8 GPUs). Contrastive losses used all-gather synchronization across ranks to ensure full batch distribution visibility for NT-Xent computation. All random number generators were seeded at 42 for reproducibility. Loss weights shared across all experiments were *λ*_con_ = 0.2, *λ*_align_ = 0.5 (except in the alignment ablation). The fitting loss was annealed linearly from 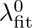 over epochs 20–100 to a floor of 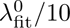.

#### A.1.2 ODE solver

All experiments used the Dormand-Prince adaptive Runge-Kutta solver (dopri5) via torchdiffeq [22] with absolute and relative tolerances of 10^−4^ and 10^−3^, respectively. Gradients were computed through the standard non-adjoint sensitivity method and the adjoint method was not used as it reduced training stability at the batch sizes required for effective contrastive learning.

### A.2 Cross-Modal Alignment Ablation

To assess how the cross-modal alignment loss shapes the trajectory-only results, we trained a model identical to Experiment (A) except *λ*_align_ = 0. Its test accuracy was 95.75%, compared with 97.40% for the full model. Without alignment, the classifier relied mainly on the time-averaged representation, and trajectory-only predictions concentrated on Working Memory and Gambling. To test whether the trajectory representation nevertheless carried condition-discriminative information, we trained a linear logistic-regression probe directly on it, with five-fold cross-validation grouped by subject. The probe reached 70.2% accuracy without alignment, compared with 77.8% for the full model, with a similar ordering across conditions, indicating that the alignment loss mainly determines whether the classifier draws on information already present in the trajectory representation.

### A.3 Null Model Testing

Trajectory-only classification accuracy was validated using label permutation, in which trajectories are held fixed while class labels are shuffled at random (1,000 permutations), directly testing whether trajectory-only accuracy exceeds chance classification independently of temporal structure. Results for Experiment A are reported in Table A2.

All conditions except Relational Reasoning significantly exceeded the label-permutation chance baseline (*p <* 0.001), indicating that trajectory-only accuracy exceeds chance classification. The high chance level for Working Memory (33.89%) reflects the tendency of the ablated classifier to assign samples from other conditions to Working Memory, so that its trajectory-only accuracy partly reflects this assignment as well as its own trajectory information. Relational Reasoning, at 0.0% with *p* = 1.0, was indistinguishable from the permutation distribution.

**Table A2.** Label permutation results.

| **Condition** | **Original (%)** | **Chance Mean (%)** | **Δ (pp)** | *p* |
|---|---|---|---|---|
| WM | 93.40 | 33.89 | +59.51 | < 0.001 |
| REST | 89.34 | 12.32 | +77.02 | < 0.001 |
| LANGUAGE | 86.29 | 10.98 | +75.31 | < 0.001 |
| SOCIAL | 83.25 | 10.76 | +72.49 | < 0.001 |
| MOTOR | 80.20 | 12.14 | +68.06 | < 0.001 |
| GAMBLING | 64.97 | 14.17 | +50.80 | < 0.001 |
| EMOTION | 28.43 | 5.40 | +23.03 | < 0.001 |
| RELATIONAL | 0.00 | 0.00 | 0.00 | 1.0 |
| Overall | 65.74 | 12.46 | +53.28 | < 0.001 |

Trajectory-only accuracy versus permutation chance mean (1,000 permutations). All *p*-values are two-sided.

### A.4 Fitting Loss Ablation

To assess how the fitting loss shapes the trajectory-only results, we trained an ablation model identical to Experiment (A) in all respects except 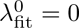, removing the fitting loss while preserving the contrastive and alignment objectives. This model reached a best validation accuracy of 96.39%, within 0.65 percentage points of the full model. Per-condition trajectory-only accuracies are reported in Table A3.

**Table A3.** Fitting loss ablation results.

| **Condition** | **Full Model (%)** | **No Fitting Loss (%)** | **Δ (pp)** |
|---|---|---|---|
| WM | 93.40 | 96.45 | +3.05 |
| REST | 89.34 | 83.76 | −5.58 |
| LANGUAGE | 86.29 | 90.86 | +4.57 |
| SOCIAL | 83.25 | 88.83 | +5.58 |
| MOTOR | 80.20 | 81.22 | +1.02 |
| GAMBLING | 64.97 | **31.47** | −33.50 |
| EMOTION | 28.43 | 36.04 | +7.61 |
| RELATIONAL | 0.00 | 0.00 | 0.00 |
| Overall | 65.74 | 63.58 | −2.16 |

The overall pattern of trajectory-only accuracy was preserved without the fitting loss, with all conditions except Gambling within 8 percentage points of the full model, indicating that the contrastive and alignment objectives largely determine the trajectory-only results. Gambling was the exception, with trajectory-only accuracy falling from 64.97% to 31.47%, indicating that the fitting loss specifically shapes trajectory geometry for this condition, which also showed the strongest within-condition relationship between trajectory path length and trajectory-only classification confidence. Together with the alignment ablation (Supplementary A3), these results indicate that the trajectory-only results depend on the training objectives, with the alignment loss determining whether the classifier draws on trajectory information and the fitting loss shaping trajectory geometry for specific conditions.

### A.5 Ergodicity and Trajectory Geometry Measures

Ergodic breakdown was operationalized through two measures computed from the extracted latent trajectories **Z**_traj_ ∈ ℝ^*N*×*T* ×*d_z_*^ and the GNN embedding sequences Z_gnn_∈ ℝ^*N* ×*T* ×*d_z_*^.

#### A.5.1 Ergodicity gap

The ergodicity gap measures the Euclidean distance between the mean of the observed embedding sequence and the mean of the trajectory generated by the learned dynamics:

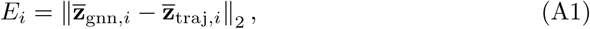

where **z̄***_i_* = *T*^−1^ Σ*_k_* **z***_i_*(*t_k_*). A smaller *E_i_* indicates that the trajectory closely tracks the embeddings, as expected when the fitting loss constrains it through a structured sequence of distinct configurations. A larger *E_i_* indicates that the trajectory mean drifts from the embedding mean, as expected when the embeddings remain near a single configuration and leave the trajectory only weakly constrained. Higher-complexity conditions (Working Memory, CARIT) therefore produce smaller gaps, and lower-complexity conditions (Relational Reasoning, Visuomotor Tracking) larger gaps. Pairwise differences between conditions were assessed using two-sided Mann–Whitney *U* tests with Cohen’s *d* as the effect size measure.

#### A.5.2 Temporal variance ratio (TVR)

The TVR compares the temporal variance of the trajectory with that of the embeddings:

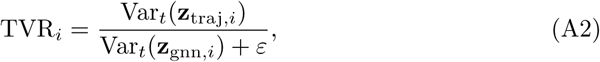

where variance is computed across time and averaged across latent dimensions, and *ε* = 10^−6^ ensures numerical stability. A TVR above 1 indicates that the trajectory varies more over time than the embeddings, which occurs most readily when the embeddings themselves vary little, as in near-stationary conditions, whereas a TVR near 1 indicates that the trajectory reproduces the temporal variance of the embeddings.

#### A.5.3 Trajectory geometry measures

Path length, tortuosity, mean curvature, mean speed, and peak excursion were computed directly from **Z**_traj_ as described in the Trajectory Feature Extractor subsection of the Methods. Dynamical richness was quantified for each condition *c* by the participation ratio,

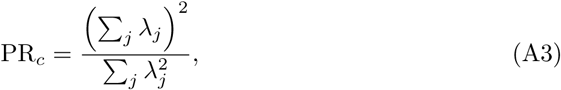

where *λ_j_* are the eigenvalues of the covariance of the trajectory states pooled across all time points and all samples of condition *c*. Differences in participation ratio between groups were assessed with 90% confidence intervals from bootstrap resampling of samples. Group comparisons of per-sample measures used two-sided Mann–Whitney *U* tests with Cohen’s *d*, and Spearman *ρ* was used for within-condition correlations between path length and trajectory-only classification confidence. All multiple comparisons were corrected using the Benjamini–Hochberg FDR procedure at *q <* 0.05.

### A.6 Behavioral Analysis

#### A.6.1 Ergodicity gap and fluid intelligence

Fluid cognition was measured with the NIH Toolbox Fluid Cognition Composite (CogFluidComp_Unadj) for HCP-YA, and fluid intelligence with FluidIQ_Tr35_60y for HCP-Aging. In HCP-Aging, the association between the resting-state ergodicity gap and fluid intelligence was assessed across subjects with Spearman correlation in Experiment (C) (*ρ* = 0.233, *p* = 0.0006, *N* = 211). Since FluidIQ_Tr35_60y is not age-adjusted and correlated with age (*ρ* = −0.537), the association was also assessed after controlling for age, by correlating the residuals of the ranks of the ergodicity gap and of fluid intelligence after linear regression of each on the ranks of age (partial Spearman correlation, *ρ* = 0.178, *p* = 0.0096).

In a complementary group analysis, subjects were median-split by fluid intelligence separately within each condition and cohort, and the ergodicity gap, path length, and temporal variance ratio were compared between groups using Mann–Whitney *U* and permutation tests (1,000 permutations), with Benjamini–Hochberg FDR correction across all tests within each cohort (24 tests for HCP-YA and 12 tests per experiment for HCP-Aging). In HCP-YA, no test survived correction in either atlas (all *q >* 0.44), and the nominally significant effects in Experiment (A), for Emotion path length (*d* = 0.302, *p* = 0.042) and Gambling temporal variance ratio (*d* = 0.259, *p* = 0.051), did not survive correction (*q* = 0.444 for both). In HCP-Aging, the only effect surviving correction was the resting-state ergodicity gap in Experiment (C) (high fluid intelligence mean = 10.77, low fluid intelligence mean = 10.23, *d* = 0.51, *p*_perm_ *<* 0.001, *q <* 0.001), and no effect survived correction in Experiment (D) (all *q >* 0.90), indicating that the association was specific to rest in older adults and to the cortical atlas.

#### A.6.2 Trajectory geometry and fluid intelligence

As an exploratory analysis, we tested whether the trajectory representation could predict fluid intelligence for individual subjects. Cross-validated prediction used a pipeline of feature standardization, principal component analysis (*n*_components_ = 5, unwhitened), and ridge regression with internally cross-validated regularization (RidgeCV), evaluated with 5-fold cross-validation and applied separately to trajectory-only features (**z**_pool_) for each condition and cohort. The number of components was chosen for the signal-to-noise regime of the data. With approximately 200 subjects per condition, using *k* components incurs an expected noise-explainable variance of *k/N* in the training folds, so that *k* = 5 gives *k/N* ≈ 0.026, limiting the fitting of noise while preserving signal. The regularization parameter was selected separately within each fold from 9 logarithmically spaced values between 10^−3^ and 10^6^, avoiding contamination between training and test data. Cross-validated *R*^2^ values were near zero or negative across all conditions and cohorts (Tables A4 and A5), with RidgeCV selecting maximum regularization in all folds, indicating that the trajectory representation does not support prediction of individual fluid intelligence. This is consistent with trajectory features being optimized for differences between conditions rather than between subjects, and does not conflict with the resting-state association above, which involves a different measure and a modest effect, explaining approximately 5% of the variance, below what cross-validated prediction can reliably detect at this sample size.

**Table A4.** Fluid intelligence prediction *R*^2^ (HCP-YA).

| **Condition** | **Cortical (Exp (A))** |  | **Cortico-subcortical (Exp (B))** |  |
|---|---|---|---|---|
|  | *R*<sup>2</sup> | **95% CI** | *R*<sup>2</sup> | **95% CI** |
| WM | −0.051 | [−0.106, −0.009] | −0.050 | [−0.114, −0.007] |
| REST | −0.057 | [−0.100, −0.026] | −0.037 | [−0.083, −0.004] |
| LANGUAGE | −0.042 | [−0.091, −0.008] | −0.009 | [−0.065, 0.042] |
| SOCIAL | −0.046 | [−0.091, −0.013] | −0.056 | [−0.105, −0.016] |
| MOTOR | −0.076 | [−0.129, −0.037] | −0.067 | [−0.131, −0.017] |
| GAMBLING | −0.044 | [−0.111, 0.010] | −0.019 | [−0.072, 0.019] |
| EMOTION | −0.034 | [−0.097, 0.011] | +0.026 | [−0.044, 0.083] |
| RELATIONAL | −0.061 | [−0.131, −0.014] | −0.079 | [−0.125, −0.050] |

Cross-validated *R*<sup>2</sup> with 95% confidence intervals from trajectory-only features. *N* = 193 (cortical) and *N* = 195 (cortico-subcortical).

**Table A5.** Fluid intelligence prediction *R*^2^ (HCP-Aging).

|  | **Cortical (Exp (C))** |  | **Cortico-subcortical (Exp (D))** |  |
|---|---|---|---|---|
| **Condition** | *R*<sup>2</sup> | **95% CI** | *R*<sup>2</sup> | **95% CI** |
| CARIT | −0.024 | [−0.070, 0.009] | −0.013 | [−0.066, 0.028] |
| REST | −0.052 | [−0.118, −0.004] | −0.038 | [−0.077, −0.014] |
| FACENAME | −0.001 | [−0.077, 0.062] | −0.020 | [−0.084, 0.028] |
| VISMOTOR | −0.030 | [−0.085, 0.010] | −0.050 | [−0.090, −0.021] |

Cross-validated *R*<sup>2</sup> with 95% confidence intervals from trajectory-only features. *N* = 211 (cortical) and *N* = 221 (cortico-subcortical).

### A.7 Aging Analysis Procedures

Aging was assessed primarily within the HCP-Aging cohort, comparing conditions under the same learned model. Ergodicity gaps were compared between CARIT and Visuomotor Tracking, the higher- and lower-complexity conditions, and between Face-Name Memory and Rest, using two-sided Mann–Whitney *U* tests with Cohen’s *d*, separately for the cortical (Experiment (C)) and cortico-subcortical (Experiment (D)) atlases. For the visualization of latent organization (Fig. 4), principal component analysis was applied to the time-averaged trajectory of each sample, separately for each atlas configuration, so that positions are comparable within but not across configurations. The association between the resting-state ergodicity gap and age within HCP-Aging was assessed with Spearman correlation across subjects in Experiment (C). Ages were taken from the cohort’s demographic records, with participants recorded as 90 or older coded as 90.

Dynamical richness was compared between cohorts for three pairs of matched conditions, Working Memory and CARIT, Relational Reasoning and Visuomotor Tracking, and Emotion and Face-Name Memory, and for Rest, which is present in both cohorts. The first two pairs are matched on chronnectome complexity, as the higher- and lower-complexity conditions of each cohort, and the third by cognitive domain. For each pair, the difference in pooled participation ratio (HCP-YA minus HCP-Aging) was computed, with 90% confidence intervals from 2,000 bootstrap resamples of samples within each condition (Table A6). These differences are reported descriptively and are not attributed to aging alone, since the two cohorts were modeled separately by models trained on different numbers of conditions.

**Table A6.** Cross-cohort comparison of dynamical richness (pooled participation ratio).

|  | **Pair** | **HCP-YA** | **HCP-Aging** | **Difference** | **90% CI** |
|---|---|---|---|---|---|
| Cortical | WM / CARIT | 6.66 | 5.76 | +0.91 | [+0.17, +1.44] |
|  | RELATIONAL / VISMOTOR | 6.35 | 5.47 | +0.88 | [+0.03, +1.53] |
|  | EMOTION / FACENAME | 8.12 | 5.72 | +2.40 | [+1.48, +2.89] |
|  | REST / REST | 9.20 | 6.35 | +2.85 | [+1.63, +3.33] |
| Cortico-subcortical | WM / CARIT | 7.66 | 5.38 | +2.29 | [+1.35, +2.66] |
|  | RELATIONAL / VISMOTOR | 7.12 | 3.84 | +3.28 | [+2.26, +3.85] |
|  | EMOTION / FACENAME | 8.95 | 4.56 | +4.39 | [+3.43, +4.70] |
|  | REST / REST | 9.03 | 5.37 | +3.66 | [+2.47, +4.23] |

Positive differences indicate higher dynamical richness in HCP-YA. Bootstrap resampling duplicates samples and thereby lowers participation ratio, the confidence intervals are asymmetric, extending further below the point estimates than above them.

**Table A7.** Sensitivity analysis of Relational Reasoning trajectory-only accuracy across loss configurations.

| **Exp.** | λ<sub>con</sub> | λ<sub>fit</sub> | λ<sub>align</sub> | **RELATIONAL (%)** |
|---|---|---|---|---|
| E112 (primary) | 0.2 | 0.05 | 0.5 | 0.00<sup>*a*</sup> |
| E112, no fit | 0.2 | 0.00 | 0.5 | 0.00<sup>*a*</sup> |
| E112, no align | 0.2 | 0.05 | 0.0 | 0.00<sup>*a*</sup> |
| E108<sup>*b*</sup> | 0.2 | 0.50 | — | 0.00<sup>*a*</sup> |
| E110 | 0.2 | 0.30 | 0.3 | 0.00<sup>*a*</sup> |
| E111 | 0.2 | 0.00 | 0.5 | 1.52<sup>*a*</sup> |
| E113 | 0.2 | 0.15 | 0.5 | 0.00<sup>*a*</sup> |
| E114 | 0.1 | 0.05 | 0.5 | 0.51<sup>*a*</sup> |

All other hyperparameters identical to Experiment (A) unless noted. <sup>*a*</sup>At or below chance (12.5%). <sup>*b*</sup>Earlier model version without the cross-modal alignment loss.

### A.8 Statistical Analysis

All statistical analyses were conducted in Python 3.10 using scipy 1.11 [55], numpy 1.24 [56], and statsmodels 0.14 [57]. Group comparisons used two-sided Mann– Whitney *U* tests with Cohen’s *d* (pooled standard deviation) as the effect size measure. Permutation tests used 1,000 iterations with random seed 42. Differences in pooled participation ratio were assessed with 90% confidence intervals from 2,000 bootstrap resamples of samples. Multiple comparisons were corrected using the Benjamini– Hochberg false discovery rate procedure [58] at *q <* 0.05, applied separately within each cohort and atlas configuration. Continuous correlations used Spearman *ρ*, and associations controlling for age used partial Spearman correlation, computed from the residuals of the ranks of each variable after linear regression on the ranks of age. Cross-validated fluid intelligence prediction used RidgeCV with 5-fold cross-validation, with R^2^ = 1 − SS_res_*/*SS_tot_ and confidence intervals from the bootstrap distribution across folds. We also assessed the sensitivity of the results to the loss configuration. Relational Reasoning trajectory-only accuracy remained at or below the eight-class chance baseline (12.5%) across all HCP-YA cortical configurations tested, varying *λ*_fit_, *λ*_con_, and *λ*_align_ (Table A7), indicating that this result is independent of the loss configuration.

### A.9 Classification Performance

Per-condition precision, recall, and F1 scores are reported for the full model classification in each experiment (Tables A8 and A9), together with the confusion matrices of the full model (Fig. A1).

**Table A8.** Per-condition precision, recall, and F1 score (%) of the full model in HCP-YA.

|  | **Cortical, Experiment (A)** |  |  | **Cortico-subcortical, Experiment (B)** |  |  |
|---|---|---|---|---|---|---|
| **Condition** | Precision | Recall | F1 | Precision | Recall | F1 |
| REST | 100.0 | 100.0 | 100.0 | 98.0 | 100.0 | 99.0 |
| EMOTION | 96.9 | 96.4 | 96.7 | 94.2 | 99.0 | 96.5 |
| GAMBLING | 97.3 | 90.4 | 93.7 | 96.3 | 80.1 | 87.5 |
| LANGUAGE | 100.0 | 99.5 | 99.7 | 99.5 | 99.0 | 99.2 |
| MOTOR | 98.5 | 100.0 | 99.2 | 98.0 | 99.0 | 98.5 |
| RELATIONAL | 92.0 | 99.5 | 95.6 | 77.0 | 97.4 | 86.0 |
| SOCIAL | 99.0 | 99.5 | 99.2 | 100.0 | 98.0 | 99.0 |
| WM | 95.9 | 93.9 | 94.9 | 95.8 | 81.1 | 87.8 |

**Table A9.** Per-condition precision, recall, and F1 score (%) of the full model in HCP-Aging.

|  | **Cortical, Experiment (C)** |  |  | **Cortico-subcortical, Experiment (D)** |  |  |
|---|---|---|---|---|---|---|
| **Condition** | Precision | Recall | F1 | Precision | Recall | F1 |
| REST | 89.6 | 95.6 | 92.5 | 79.8 | 98.8 | 88.3 |
| CARIT | 91.3 | 84.4 | 87.8 | 96.1 | 82.5 | 88.8 |
| FACENAME | 90.9 | 93.8 | 92.3 | 98.6 | 90.8 | 94.6 |
| VISMOTOR | 97.3 | 95.1 | 96.2 | 98.7 | 97.1 | 97.9 |

**Fig. A1.**
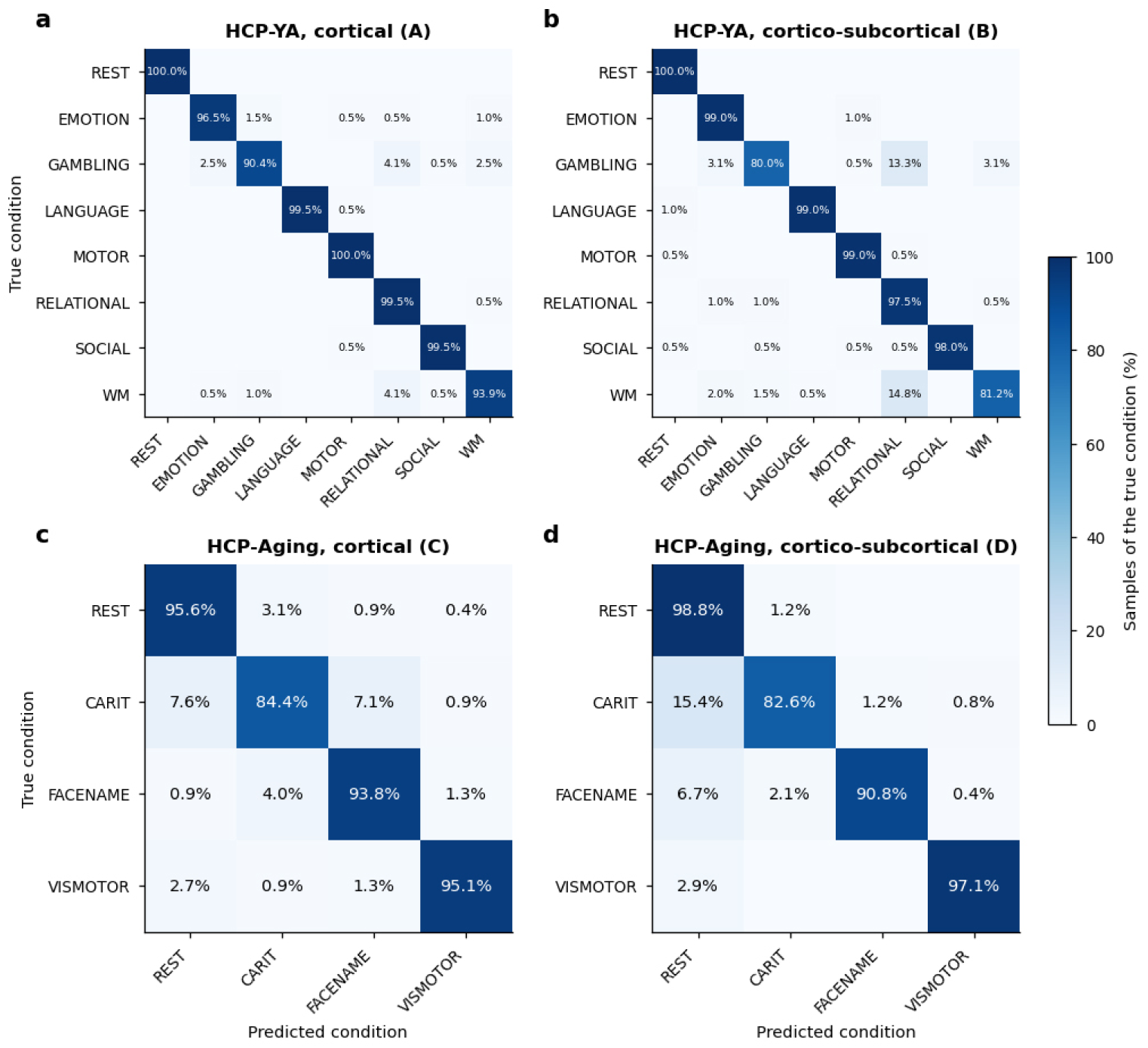
Confusion matrices of the full model in each experiment. (**a**) HCP-YA, cortical atlas, Experiment (A). (**b**) HCP-YA, cortico-subcortical atlas, Experiment (B). (**c**) HCP-Aging, cortical atlas, Experiment (C). (**d**) HCP-Aging, cortico-subcortical atlas, Experiment (D). Rows indicate true conditions and columns predicted conditions. Matrices are normalized by row, so that each entry gives the percentage of a true condition’s samples assigned to each predicted condition and diagonal entries give the recall of each condition, as reported in Supplementary Tables A8 and A9. Values are rounded to one decimal place.

